# Neural correlates of co-players’ identity and reputation processing on cooperation in the public goods game

**DOI:** 10.1101/2024.10.30.620899

**Authors:** Waldir M. Sampaio, Fernanda Naomi Pantaleão, Beatriz de Souza, Ana Luisa Freitas, Paulo S. Boggio

## Abstract

This study investigates the neural correlates of identity and reputation processing in cooperative decision-making during the Public Goods Game (PGG). Using functional near-infrared spectroscopy (fNIRS), we examined how different types of social information—co-player facial photos, reputational scores, and a combination of both—influence activity in the dorsolateral prefrontal cortex (dlPFC) and ventromedial prefrontal cortex (vmPFC). Our findings suggest that the dlPFC is crucial for integrating reputational information, reflecting its role in cognitive control and modulation of expectations during cooperative interactions. The vmPFC, on the other hand, is involved in attributing value to the cooperative decision by synthesizing identity and reputation cues. Reputation alone elicited greater dlPFC activity, highlighting the cognitive demands of evaluating past behaviors. When both identity and reputation information were presented, vmPFC activity increased, indicating greater value processing. These results demonstrate the distinct but complementary roles of the dlPFC and vmPFC in processing social cues during cooperative decision-making.

---

Cooperation is defined as the interdependent action of two or more people towards the production of mutual goods (Dovidio et al., 2006) and, as such, it is a behaviour that depends on gathering information about the individuals with whom one is interacting. Identifying the people with whom you are cooperating is crucial, as indirect reciprocity relies on the recognition of the co-players’ and the assessment of their reputations (Nowak & Sigmund, 1998). Understanding how individuals evaluate the potential for cooperation and respond to this information is central to understanding prosocial behaviour.

Economic games, such as the Public Goods Game (PGG), are widely used to study cooperation (Feygina & Henry, 2015). In a typical PGG game, three or more players receive an initial amount of money and must decide whether to donate this sum to a common fund or to keep it to themselves. After all decisions are made, the total amount donated to the fund is doubled and equally distributed among all group members, including those who did not donate initially (referred to as free-riders). Consequently, free-riders end up with a higher value than the members who contributed to the common fund. In this context, cooperative behaviour is defined by the act of contributing to the group. This setup allows for experimental manipulations designed to explore the effects of various factors that can influence prosocial behaviour.

Previous studies using the PGG have demonstrated that context can modulate cooperation. For example, revealing information about co-players, such as their identity, face photos or reputation (based on their previous game behaviour) - influence cooperative behaviour, shaping participants’ expectations as they evaluate how likely others are to reciprocate their prosocial actions (Andreoni & Petrie, 2004; Christens et al., 2019; Milinski et al., 2002; Sylwester & Roberts, 2010).

Our previous investigation replicated the finding that revealing identities through photos of co-players in a PGG, along with their reputations, influenced cooperative behaviours. Specifically, showing faces did not directly increase cooperation but, when associated with a co-player reputation, it modulated it. Reputation alone significantly affected cooperation, as it decreased with free-rider reputations and increased with cooperative reputations. When photos were presented alongside reputation information, the faces mitigated the negative effects of free-rider reputations and enhanced cooperation for both neutral and cooperative reputations (Sampaio et al., 2023). Building on these findings, our ongoing work demonstrates that the effect does not appear to be associated with the modulation of reputation based on social categories such as age, gender, and race, as homophily in these dimensions did not influence reputation perception across three different studies (Sampaio et al., 2024). However, in the same investigation, it was found that race influenced the cooperative behaviours of White participants, who cooperated more with similar cooperative co-players in comparison with Black participants.

There is a possibility that these results could be explained by the neurophysiological processing of different types of information. Literature has shown that both ventromedial (vmPFC) and dorsolateral prefrontal cortices (dlPFC) play important roles in cooperative decision-making processes (Hackel et al., 2020; Pärnamets et al., 2020; Wills et al., 2018). The dlPFC is considered responsible for modulating the value of cooperation decisions by integrating information about goals, individual motivations, and modulation of expectations (Pärnamets et al., 2020; Tusche & Hutcherson, 2018). In contrast, the vmPFC is considered an integrational hub that attributes value to the options under consideration in cooperative decisions (Cutler & Campbell-Meiklejohn, 2019; Levy & Glimcher, 2012; Ruff & Fehr, 2014).

## Our research

Our research aims to explore how different types of information about co-players in cooperative decision-making are processed in the cortex, with a particular focus on the roles of the ventromedial prefrontal cortex (vmPFC) and dorsolateral prefrontal cortex (dlPFC). In our experiment, we present participants with three types of information about their co-players: facial photos, reputational scores based on past behaviour, and a combination of both. This approach builds on previous research that has examined the effects of identity and reputation in public goods games, notably the study by Sampaio et al. (Sampaio, 2023), which highlights the importance of understanding how these variables interact.

The dlPFC is thought to play a key role in weighing the value of cooperative decisions by integrating goal-relevant information and managing motivations and expectations during social interactions (Carlson & Crockett, 2018; Tusche & Hutcherson, 2018). Research indicates that the dlPFC is relevant in strategic decision-making (Wout et al., 2005) and self-control in social interactions (Rilling & Sanfey, 2009). We expect that the kind of information provided—whether identity-related, reputational, or both—will modulate neural activity in the dlPFC; specifically, reputational scores, which are closely tied to past actions and therefore modulate expectations and may elicit self-control, are likely to elicit greater dlPFC activation compared to mere facial identity. This aligns with findings that suggest the dlPFC is involved in processing moral dilemmas and deliberate decision-making (Hare et al., 2009).

On the other hand, we hypothesise that the vmPFC, known for its role in attributing value and processing rewards, might be more active when participants receive personal information about their co-players, which enhances the perceived value of cooperating. Studies have shown that the vmPFC is engaged when individuals evaluate financially rewarding stimuli (Van Overwalle, 2009) and when they evaluate social stimuli with personal relevance (Northoff et al., 2011). Furthermore, the vmPFC has been shown to be sensitive to available knowledge about others (Dang et al., 2019). Therefore, when both identity and reputation information are available, we anticipate that this integrated valuation will be reflected in increased vmPFC activity, as participants synthesise multiple cues to make a well-rounded judgement about their co-players. This suggests that the vmPFC may facilitate decision-making by providing a valuation framework that incorporates both identity and reputation, thereby influencing prosocial behaviour in social dilemmas.

Using functional near-infrared spectroscopy (fNIRS) alongside the Public Goods Game (PGG), our study aims to understand how these prefrontal regions contribute to the processing of cooperative decision-making depending on the type of interpersonal information provided. It’s important to highlight that our goals and hypotheses are centred on understanding the role of the vmPFC and dlPFC in processing different types of social information available in a cooperative setting, rather than focusing on the direct impact of these brain regions on a specific outcome (i.e. cooperative or free-rider response). Our interest is to explore how identity and reputation cues are integrated by these prefrontal regions during decision-making.

Therefore, this study aimed to investigate the neural processes underlying cooperation through the modulation of identity and reputation of co-players and their effects on prosocial behaviour, using Functional Near-Infrared Spectroscopy (fNIRS) and the PGG. We hypothesised that variables influencing participants’ expectations about other players would increase dlPFC activity, but reputation indices should elicit more activation than photos of faces, since reputation seems to be more closely tied to expectations than photos. Regarding vmPFC, we expected greater activity when participants receive more personal information about their co-players, due to its involvement in the valuation system and in the process of social information.

## 5.1 Methods

### 5.1.1 Participants

A total of 162 participants volunteered to take part in the experiment, which involved playing the PGG during fNIRS data collection. 7 participants were excluded due to invalid responses (over 6 empty values) and invariant responses (consistently responded with 0 or 1 in all trials), resulting in a total of 155 participants (average age = 23.03, SD = 5.14, 128 female, 31 male, 4 did not respond). They were randomly assigned to one of four experimental groups, which varied in the information they received about other players during the PGG: (i) control (no identity or reputation information, (ii) only reputation (neutral, free-rider, or cooperative), (iii) only face, and (iv) face and reputation (neutral, free-rider, or cooperative).

Information about the study was shared in the laboratory’s communication channels and social media platforms. All participants were informed that they could withdraw their consent from the experiment at any time, and each provided informed consent. The entire experiment was approved by the Institutional Ethics Committee of Mackenzie University (SISNEP, Brazil; CAAE: 46235921.0.0000.0084) and adhered strictly to the guidelines of the Declaration of Helsinki.

### 5.1.2 Instruments

#### Public goods game

Similarly to Sampaio et al. (2023), we utilised a four-player one-shot version of the Public Goods Game (PGG), closely resembling the design employed by Hackel and colleagues (2020), and adapted it for online use with the PsyToolkit tool, with the co-players programmed by the researchers. Prior to gameplay, participants viewed an instructional screen informing them that they were not interacting with real-time players and that their contributions in each round had been recorded beforehand. They were also made aware that their responses would be recorded, making it clear that their input could potentially contribute to future research with the following statement: “Your responses will be recorded anonymously and utilised in future studies.” The game comprised 20 rounds, with participants engaging with three unknown players who appeared only once throughout the game. Participants were not told how many rounds they would play before starting. In each round, they received a fictitious amount of R$8.00, which they could either retain or contribute any amount to a “public fund.”. It was explained that each player’s contribution would be doubled and evenly distributed among all four participants. After each round, they received feedback screens displaying the choices made by other players, the total amount collected in the fund, and the distribution of the public fund. Each trial consisted of four screens: an initial screen displaying player identities, where participants decided to keep or give, followed by three feedback screens showing the decisions of other players (keep or give), the total fund amount, and its distribution among players, each displayed for 3 seconds. The sequence of decisions made by co-players was consistent across all groups, ensuring that any order effects were identical for all participants.

For this version, the four experimental groups played the same version of the PGG, with the only variation being the information they received about the other players at the beginning of the rounds. This information included the players’ identities (photo) and/or a group reputation indicating how often the three players had cooperated in the previous 20 rounds. The reputation classifications included were presented with bar graphs on the screen and could vary as follows: free-rider (≤ 25% cooperation in the last 20 rounds), neutral (≥ 50% and < 75% cooperation), and cooperative (≥ 75% cooperation).

The other players were not real individuals; their decisions were pre-programmed by the researcher, and the photos used did not belong to actual people but were generated by Generative Adversarial Networks (www.thispersondoesnotexist.com) to prevent any recognition by participants. Throughout the 20 rounds, all participants encountered the same player decisions. On average, the other players contributed to the public fund 59% of the time. To minimise bias towards specific racial groups, we validated photos based on perceived skin colour, selecting only those identified as white. Finally, participants received a debriefing only after the experiment concluded.

### 5.1.3 Procedure

Participants were recruited through social media advertising. After scheduling a session, participants were presented with the instructions for the experiment, as well as an informed consent form explaining other details concerning the experiment, such as benefits, risks, anonymity, data storage, and the contact information of the responsible researcher. In case of agreement with the informed consent, they answered demographic questions about sex (male, female or other), age (in years), their personal income (9 levels: R$1–R$500; R$501–R$1,000; R$1,001–R$2,000; 2,001–R$ 3,000; R$ 3,001–R$5,000; $5,001–R$10,000; R$10,001–R$20,000; R$20,001–R$100,000; R$100,001 and above), and racial identification (white; black; asian; mixed-race (pardo); native Brazilian indigenous). Then, participants were prepared for fNIRS recording, as they were explained how to play the PGG. After fNIRS cap setup, participants played the PGG. In the end, they were debriefed about the other players.

### 5.1.4 Neurophysiological data collection and probe design

Cerebral hemodynamics were measured using fNIRS (Brainsight, Rogue Resolutions LTD., Cardiff, Canada). Infrared wavelengths of 685 and 830 nm were applied at a power of 20 mW. fNIRS data were sampled at 10 Hz using the Brainsight software (Brainsight, Rogue Research, Quebec, Canada). Sources and detectors were positioned to capture cortical activity in the left and right dorsolateral prefrontal cortex (dlPFC) (Sanches et al., 2009; Zimeo Morais et al., 2018) and the ventromedial prefrontal cortex (vmPFC) (Finger et al., 2008) (see Table 1). These areas were selected due to their involvement in modulating the value of cooperation decisions, integrating goals and motivations (dlPFC) and valuation system and subjective values in decision-making processes (vmPFC) (Pärnamets et al., 2020). A total of 12 channels (4 sources and 9 detectors) were used to record activity in the target regions, with an interoptode distance ranging from approximately 24 mm to 30 mm; 4 short-separation channels (∼15 mm from the source) were also included to estimate systemic responses in the fNIRS data. Specifically, sources were placed at the following locations in the International 10-10 System: AF3, AF4, FC5, FC6; detectors were placed at: Fpz, Fp1, Fp2, F4, FT8, AF8, F3, FT7, AF7; and short-separation optodes were positioned between the following locations: AF4 and Fpz, AF3 and Fpz, FC6 and AF8, FC5 and AF7.

**Table 1.**
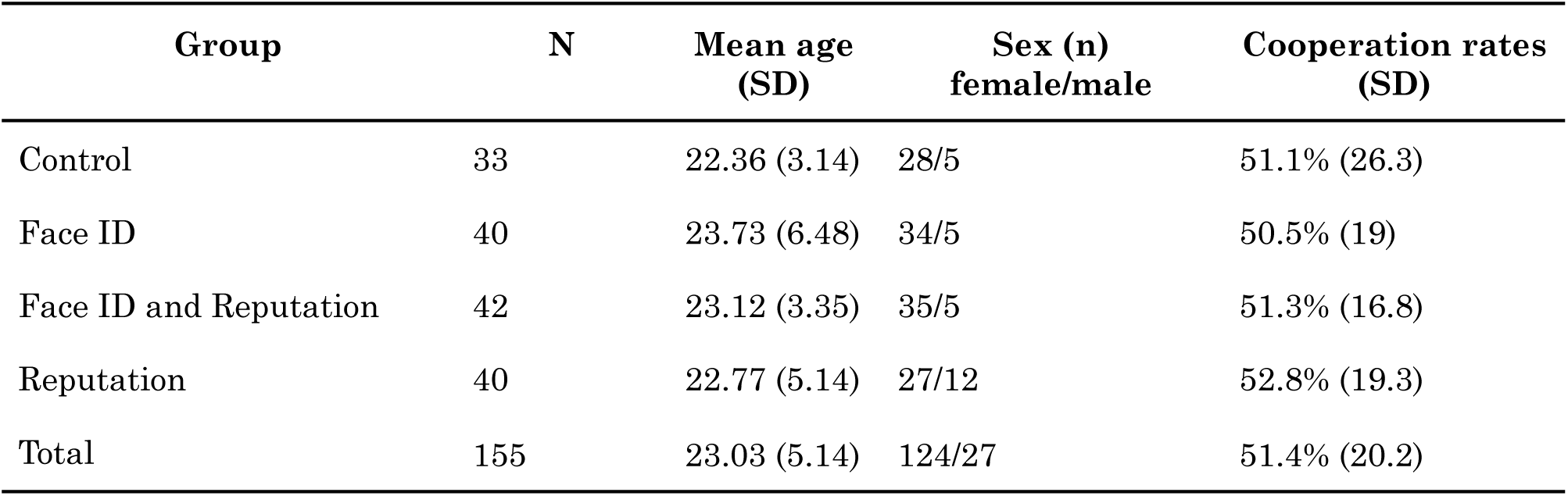
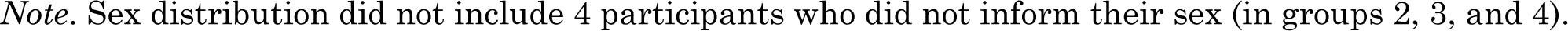
Sample size, mean age (in years), sex, and mean cooperation rates in each experimental group.

| Group | N | Mean age (SD) | Sex (n) female/male | Cooperation rates (SD) |
| --- | --- | --- | --- | --- |
| Control | 33 | 22.36 (3.14) | 28/5 | 51.1% (26.3) |
| Face ID | 40 | 23.73 (6.48) | 34/5 | 50.5% (19) |
| Face ID and Reputation | 42 | 23.12 (3.35) | 35/5 | 51.3% (16.8) |
| Reputation | 40 | 22.77 (5.14) | 27/12 | 52.8% (19.3) |
| Total | 155 | 23.03 (5.14) | 124/27 | 51.4% (20.2) |

*Note.* Sex distribution did not include 4 participants who did not inform their sex (in groups 2, 3, and 4).

### 5.1.5 Statistical analysis

#### Behavioural Results

We recorded participants’ responses—whether they cooperated with the public fund in each round (variable ‘Response’; levels: 0 = Free-ride and 1 = Cooperate)—along with the reaction time (RT) for each proposal. Responses with extreme RTs (i.e., shorter than 200 ms or longer than 20 s) were marked as null, as these might suggest inattentiveness to the task. Following this, we applied a log transformation to the RTs and identified outliers using the interquartile range (IQR) rule, which flags values 1.5 times beyond the IQR from the first and third quartiles, marking those responses as null. Participants who had more than 20% null responses were excluded from further analysis. Additionally, participants who consistently responded the same way across all rounds (i.e., always cooperating or always derogating) were also excluded, as this behaviour could reflect insensitivity to the experimental manipulations. The exclusion of invariant responses was based on both theoretical and methodological considerations. Our experimental design, particularly the reputation indices, was intended to prompt variability in cooperative behaviours, and participants whose responses remained unchanged across different conditions may not have engaged meaningfully with the stimuli. Moreover, in economic games like the ones in this study, some invariant response patterns may be optimal, but excluding them is warranted since our goal is to explore behavioural variability in response to changing social cues and dynamics. Invariant responses could also indicate lack of engagement in the task, which is a major concern in online studies where direct control over participant behaviour is limited.

For the main analysis, we used a logistic mixed model with the ‘glmer’ function from the lme4 package, employing maximum-likelihood estimation and the ‘BOBYQA’ optimizer to predict the ‘Response’ variable (i.e., decision to cooperate in the PGG) based on the ‘Condition’ factor. The ‘Condition’ factor was derived from the four experimental groups, with those assessing reputation further broken down into three conditions (free-rider, neutral, and cooperative), creating a total of 7 levels: (0) Control (reference group in the model); (1) reputation only—free-rider; (2) reputation only—neutral; (3) reputation only—cooperative; (4) face only; (5) face and reputation—free-rider; (6) face and reputation—neutral; (7) face and reputation—cooperative. To account for random effects, the model included participant ID nested within the Group as a random effect. The formula for the main model was ‘Response∼Conditions(1|ID/Group)’. We compared the main model with alternative models that included additional predictors, but no significant differences were found. Standardised parameters were obtained by fitting the model to a standardised version of the dataset. Confidence intervals (95% CIs) and p-values were calculated using the Wald approximation. All analyses were conducted using R and R Studio.

#### Neurophysiological Results

All fNIRS data were processed using Homer 3 (version 1.80.2) (Huppert et al., 2009) and extracted with MATLAB (version R2020a) following the processing stream outlined by Yücel et. al (2021): the function **hmrR_PruneChannels** (dRange: 0 1e+07, SNRthresh: 4, SDrange: 0.0 45.0) was applied to remove channels with means outside the specified dRange, a signal-to-noise ratio (SNR) greater than SNRthresh, or a source-detector distance exceeding 45 mm. Then, **hmrR_Intensity2OD** was used to convert intensity data to optical density, followed by **hmrR_MotionArtifactByChannel** (tMotion: 0.5, tMask: 1.0, STDEVthresh: 20.0, AMPthresh: 0.20) to detect motion artefacts. Motion correction was performed with **hmrR_MotionCorrect** (iqr: 1.50) using a wavelet transformation. To enhance the signal-to-noise ratio, **hmrR_BandpassFilt** (hpf: 0.0, lpf: 0.090) was applied, and **hmrR_OD2Conc** (ppf: 1.0 1.0) was used to convert optical density data into concentration. Finally, **hmrR_BlockAvg** was employed to estimate hemodynamic responses.

We extracted the HRF values from the Homer output structure, which were exported for further analysis in R. We included data from all channels for all analysis. For all reported analysis HbO values were then multiplied by a factor of 100,000 to improve visual interpretation and maintain consistency across different scales.

Then, we calculated an ANOVA with the between-subjects factor “Group” (with 4 levels: Control, Face ID, Reputation, and Face ID and Reputation), with oxyhemoglobin during decision-making as our dependent variable. This analysis allowed for the identification of significant differences in HbO levels between the experimental conditions and across different brain regions.

Post-hoc pairwise comparisons were performed to further investigate specific differences between the groups. These comparisons focused on identifying which groups differed significantly in terms of HbO values, with results presented in the form of p-values, t-statistics, and effect sizes (Cohen’s d). The function used for post-hoc analysis was ‘pairwise.t.test’, without correction for multiple comparisons, as our study follows a hypothesis-driven approach with predefined comparisons. Given our specific hypotheses and focus on targeted effects, applying corrections for multiple comparisons was deemed unnecessary to avoid overly conservative results that might obscure meaningful findings.

The statistical t-values from our post-hoc analyses were transformed into image files using the xjview toolbox. Following Binnquist’s (2023/2024) automated script, these images were then projected onto a three-dimensional brain surface model using Surf Ice 6.

## 5.2 Results

### 5.2.1 Behavioural Results

All 162 participants completed the experiment. We excluded 7 participants due to null or invariant responses, meaning they consistently chose to either free-ride or cooperate. Demographic information for the remaining 155 participants in the analysis is presented in Table 8, along with the average cooperation rates for each group. We tested differences among the six experimental groups on sociodemographic characteristics, such as age [F(3,3336) = 7.29, p < .001] and sex [χ2(3) = 69.49, p < .001], which were both significant.

#### Behaviour on PGG

Overall, participants tended to cooperate in the PGG (cooperation rate mean: 51.4%; SD: 20.2). Using the cooperation rate values for each participant, we conducted a one-way ANOVA to compare cooperation rates across the experimental groups, which indicated no main effect for Group [F(3,151) = 0.08, p = 0.96]. Participants’ cooperation rate in the first versus in the last round did not vary significantly [χ2 = 0.85, *p* = 0.36].

#### Predicting cooperation from condition

We ran a logistic mixed model to predict Cooperation from the Condition variable, with the control group as reference. Regarding the random effects, the Group:ID variance is 0.13 (SD: 0.36), and ID variance is 0.58 (SD: 0.76). Coefficient estimates, confidence interval, and p-values of the model are described in Table 2.

**Table 2.** Logistic mixed model estimates and standard error (SE) for each term, with 95% confidence interval and *p*-value. ***0.001; **0.01; *0.05.

| Term | Estimate | SE | 95% CI | *p*-value |
| --- | --- | --- | --- | --- |
| Intercept | 0.05928 | 0.17105 | -0.28, 0.40 | 0.73 |
| Face ID | -0.04071 | 0.22902 | -0.49, 0.41 | 0.86 |
| Reputation Free-rider | -0.50264 | 0.25134 | -1.00, -0.01 | 0.05* |
| Reputation Neutral | -0.01293 | 0.25500 | -0.51, 0.49 | 0.96 |
| Reputation Cooperative | 0.69924 | 0.25363 | 0.20, 1.20 | 0.01** |
| Face ID and Rep. Free-rider | -1.23284 | 0.25284 | -1.73, -0.74 | < .001 |
| Face ID and Rep. Neutral | 0.14229 | 0.24930 | -0.35, 0.63 | 0.57 |
| Face ID and Rep. Cooperative | 1.14376 | 0.25366 | 0.65, 1.64 | < .001 |

*Note.*\*\*\*0.001; \*\*0.01; \*0.05.

The model showed that, compared to the control group, the ‘only reputation’ group significantly decreased cooperation in the free-rider condition and increased cooperation during cooperative conditions. Additionally, the ‘face and reputation’ group significantly increased cooperation during neutral and cooperative conditions. Those detected effects and their odds ratios can be seen in Figure 2.

**Figure 1.**
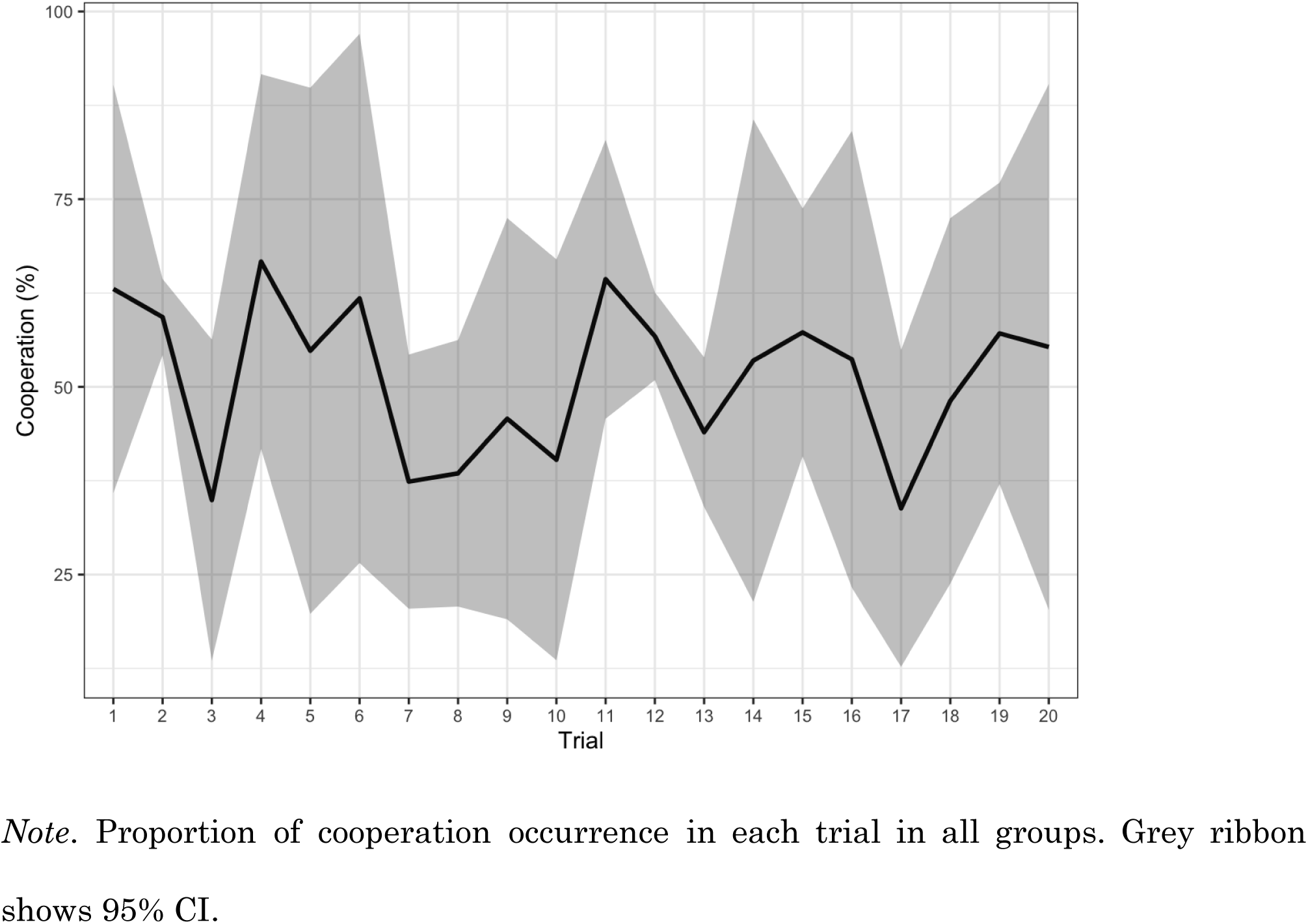
Cooperation occurrence along trials.

**Figure 2.**
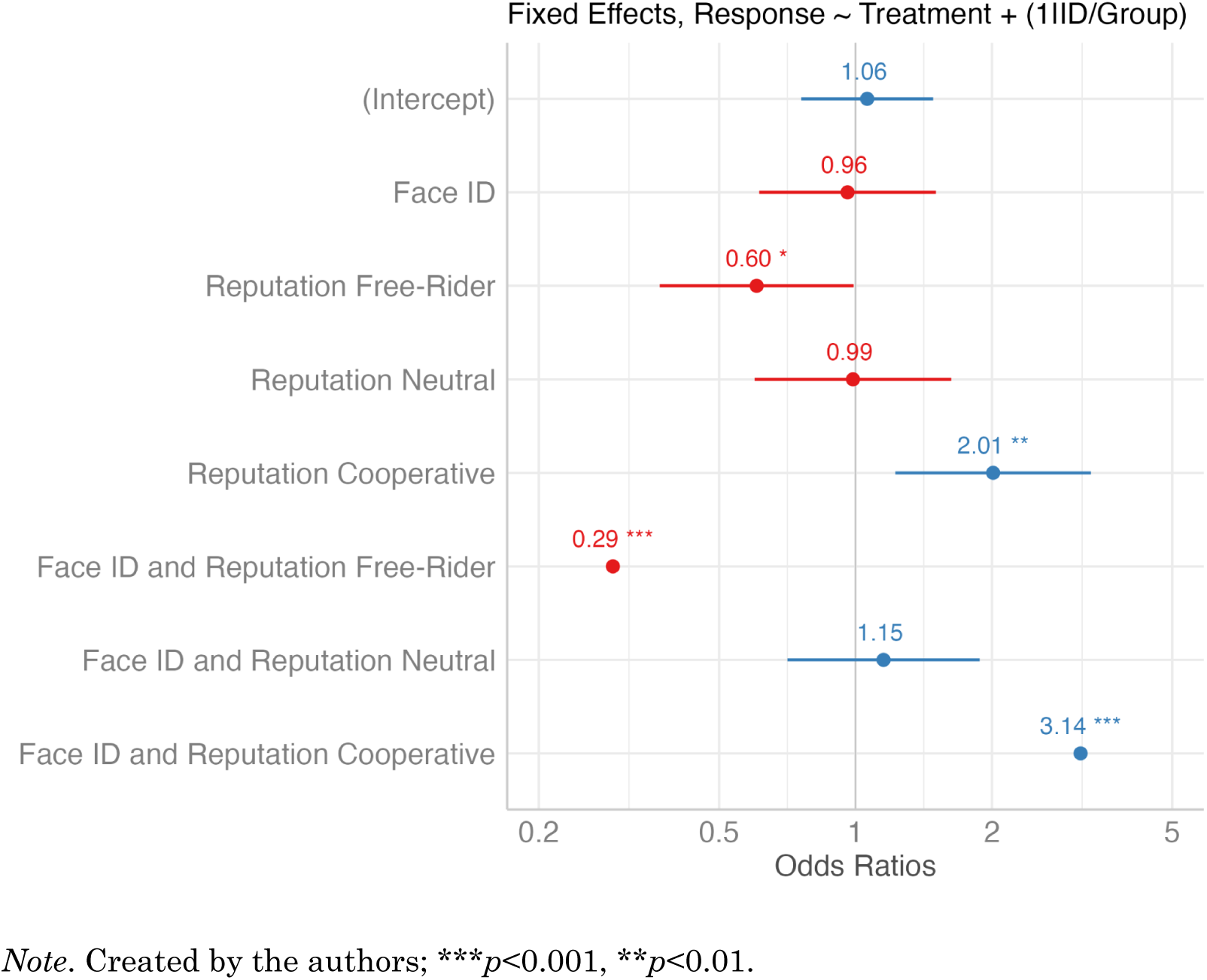
Odds ratios for the fixed effects.

### 5.2.2 Neurophysiological Results

A one-way ANOVA was conducted with the between-subjects factor “Group” (Control, Face ID, Reputation, Face ID and Reputation) and with HbO during decision-making as the dependent variable. The ANOVA revealed a significant main effect of Group on HbO levels, F(3, 5074) = 7.12, p < .001, indicating significant differences in HbO responses across the experimental conditions. The main effect of Channel, F(15, 5074) = 1.28, p = .207, and the Group x Channel interaction, F(45, 5074) = 1.10, p = .293, were not significant.

Post-hoc pairwise comparisons showed that the Face ID and Reputation group had significantly higher HbO values compared to the Control group, p = .003, and the Face ID group, p = .001. The Reputation group also had significantly higher HbO values compared to the Control group, p = .001, and the Face ID group, p < .001. There were no significant differences between the Face ID and Reputation groups, p = .721.

Analyses of specific channels further supported these findings. In Channel 1,3 (Left dlPFC), the Face ID and Reputation group had significantly higher HbO levels compared to the Control group, p = .002, the Face ID group, p < .001, and the Reputation group, p = .008. Similar patterns were observed in Channels 2,4 and 3,8, with the Face ID and Reputation group showing significantly higher values compared to other conditions. Detailed results for these comparisons are presented in Table 3 and post hoc activation t-values are presented in Figure 3.

**Figure 3.**
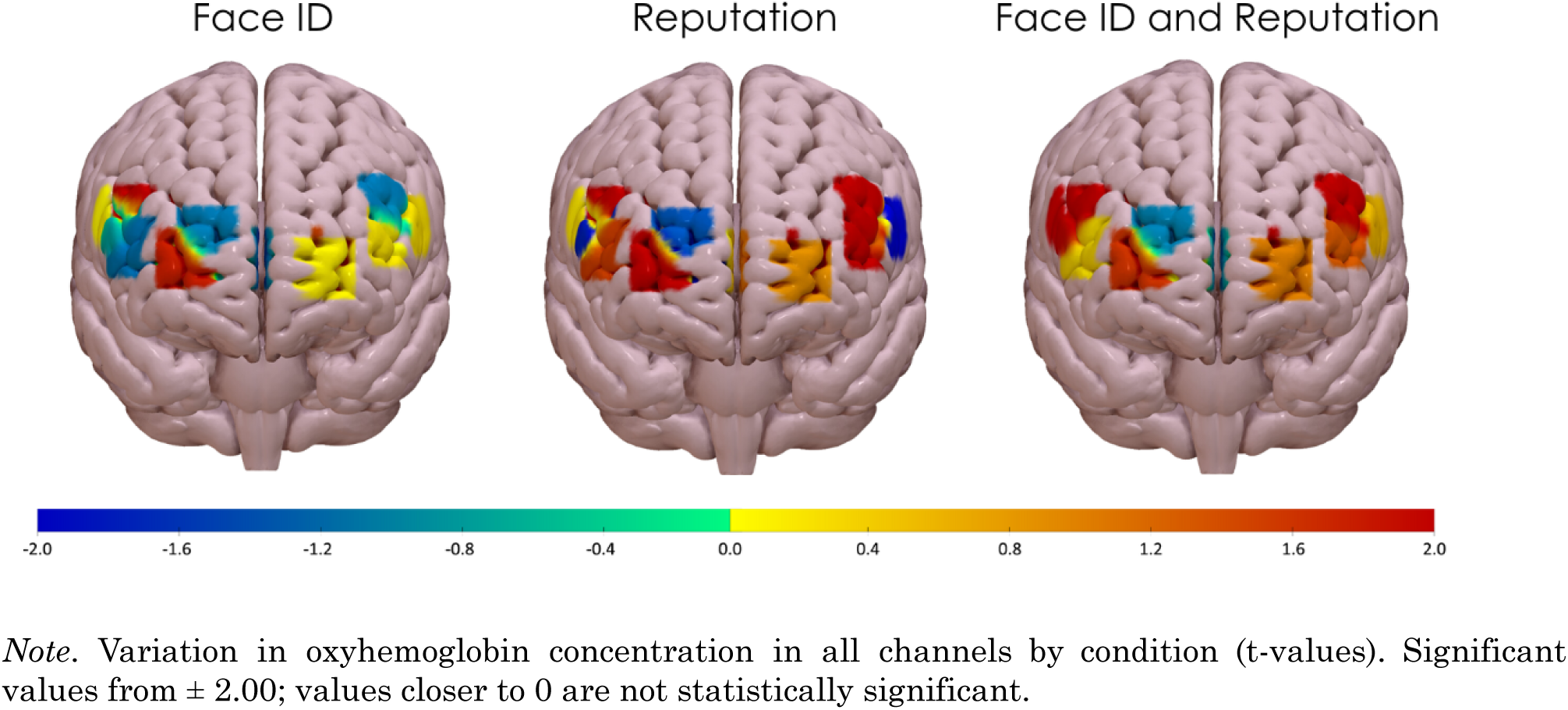
Hemodynamic response of Face ID, Reputation, and Face ID and Reputation groups compared to baseline.

**Table 3.** List of significant comparisons between the same Source-Detector between groups.

| Source -<br>Detector | ROI | Comparison | t-Value | p-Value | Cohen's<br>d | Mean<br>G1 | SD<br>G1 | Mean<br>G2 | SD<br>G2 |
| --- | --- | --- | --- | --- | --- | --- | --- | --- | --- |
| 1,3 | Left dlPFC | Control vs Face ID and Reputation | 3.1775 | .0018 | 0.4172 | 0.1766 | 2.1634 | -1.4808 | 4.4073 |
| 1,3 | Left dlPFC | Control vs Reputation | 2.2324 | .0274 | 0.3161 | 0.1766 | 2.1634 | -0.8998 | 3.7366 |
| 1,3 | Left dlPFC | Face ID and Reputation vs Face ID | -3.4550 | .0009 | -0.5629 | -1.4808 | 4.4073 | 0.8953 | 3.5940 |
| 1,3 | Left dlPFC | Face ID vs Reputation | 2.7267 | .0080 | 0.4850 | 0.8953 | 3.5940 | -0.8998 | 3.7366 |
| 1,4 | Left dlPFC | Control vs Reputation | 2.0245 | .0450 | 0.2808 | -0.0198 | 3.0185 | -1.4211 | 5.5056 |
| 2,1 | Left vmPFC | Face ID and Reputation vs Reputation | 3.8241 | .0002 | 0.4957 | 0.0636 | 2.9946 | -1.6807 | 3.9934 |
| 2,1 | Left vmPFC | Face ID vs Reputation | 3.0362 | .0032 | 0.5144 | 0.3090 | 3.4736 | -1.6807 | 3.9934 |
| 2,4 | Left vmPFC | Control vs Reputation | 4.5536 | .0000 | 0.6350 | 0.1211 | 1.6192 | -1.5947 | 2.9701 |
| 2,4 | Left vmPFC | Face ID and Reputation vs Reputation | 3.8924 | .0001 | 0.5026 | -0.1708 | 2.6953 | -1.5947 | 2.9701 |
| 2,4 | Left vmPFC | Face ID vs Reputation | 2.6917 | .0084 | 0.4138 | -0.4476 | 2.0675 | -1.5947 | 2.9701 |
| 2,13 | Left vmPFC | Control vs Face ID and Reputation | 2.6096 | .0100 | 0.3362 | -0.1968 | 0.8882 | -0.7747 | 1.9144 |
| 2,13 | Left vmPFC | Control vs Reputation | 2.4957 | .0138 | 0.2759 | -0.1968 | 0.8882 | -1.4314 | 5.1582 |
| 3,6 | Right dlPFC | Face ID and Reputation vs Face ID | -2.7943 | .0063 | -0.4248 | -3.0709 | 8.2506 | 0.2085 | 5.8122 |
| 3,6 | Right dlPFC | Face ID and Reputation vs Reputation | -2.2983 | .0224 | -0.2969 | -3.0709 | 8.2506 | -0.4559 | 9.3523 |
| 3,7 | Right dlPFC | Control vs Face ID and Reputation | 2.1049 | .0387 | 0.3658 | 0.2265 | 3.2383 | -1.0269 | 3.4862 |
| 3,8 | Right vmPFC | Control vs Face ID and Reputation | 3.2188 | .0018 | 0.5199 | 1.1838 | 3.5561 | -1.0197 | 4.4391 |
| 3,8 | Right vmPFC | Control vs Face ID | 2.0605 | .0428 | 0.4551 | 1.1838 | 3.5561 | -0.6466 | 4.4395 |
| 3,8 | Right vmPFC | Control vs Reputation | 3.0079 | .0032 | 0.4346 | 1.1838 | 3.5561 | -1.1404 | 5.8356 |
| 3,10 | Right vmPFC | Face ID vs Reputation | 2.6279 | .0099 | 0.3959 | 0.4283 | 3.4118 | -1.4324 | 5.0653 |
| 4,9 | Right dlPFC | Face ID and Reputation vs Face ID | -2.0219 | .0449 | -0.2479 | -2.4244 | 11.5650 | 0.0884 | 3.9018 |
| 4,9 | Right dlPFC | Face ID vs Reputation | 2.0079 | .0471 | 0.2979 | 0.0884 | 3.9018 | -1.5875 | 6.1160 |
| 4,11 | Right dlPFC | Control vs Reputation | 2.2340 | .0277 | 0.3461 | 1.1315 | 4.6696 | -1.0009 | 6.6042 |

Note. ROI = Region of interest; G1 = first comparison group; G2 = second comparison group.

In the dlPFC, the Control group demonstrated a decrease in baseline oxyhaemoglobin concentration compared to other groups, suggesting reduced activation. The Reputation group showed increased oxyhaemoglobin concentration, indicating heightened cognitive engagement compared to baseline. The combined Face ID and Reputation group exhibited even higher oxyhaemoglobin levels, reflecting the joint influence of identity and reputation, while the Face ID group alone showed an increase, though less pronounced compared to the combined condition. Interestingly, the Right dlPFC showed a decrease in oxyhaemoglobin concentration in the combined Face ID and Reputation group compared to the Face ID group, possibly reflecting an attenuation of cognitive load (see Table 4).

**Table 4.** Significant Effects - dlPFC (Dorsolateral Prefrontal Cortex)

| Group | Effect |
| --- | --- |
| Left dlPFC - Control | ↓ Baseline Oxyhaemoglobin concentration compared to other groups |
| Left dlPFC - Face ID | ↑ Oxyhaemoglobin concentration, but reduced when compared with combined group (Face ID and Reputation) |
| Left dlPFC - Reputation | ↑ Oxyhaemoglobin concentration, indicating increased cognitive engagement compared to baseline |
| Left dlPFC - Face ID and Reputation | ↑ Oxyhaemoglobin concentration, showing combined identity and reputation effects |
| Right dlPFC - Face ID and Reputation | ↓ Oxyhaemoglobin concentration when compared to Face ID alone, possibly reflecting attenuation of cognitive load |

*Note.* Hemodynamic responses in the dorsolateral prefrontal cortex (dlPFC) across different groups. The table summarises the effects of group conditions on oxyhaemoglobin concentration.

In the vmPFC, the Control group exhibited increased oxyhaemoglobin concentration, indicating the absence of modulatory effects. The Reputation group showed a decrease in oxyhaemoglobin concentration, suggesting valuation effects. The Face ID and Reputation group demonstrated an increase in concentration, influenced by the presence of multiple sources of social information. The Right vmPFC also showed differential effects: the Control group had increased baseline concentrations in the absence of social cues, while both the Reputation and Face ID and Reputation groups exhibited decreased oxyhaemoglobin concentrations, indicating reduced valuation in more complex social contexts (see Table 5).

**Table 5.** Significant Effects - vmPFC (Ventromedial Prefrontal Cortex)

| Group | Effect |
| --- | --- |
| Left vmPFC - Control | ↑ Oxyhaemoglobin concentration, no modulatory effects present |
| Left vmPFC - Reputation | ↓ Oxyhaemoglobin concentration, indicating valuation effects compared to baseline |
| Left vmPFC - Face ID and Reputation | ↑ Oxyhaemoglobin concentration, influenced by multi-source information |
| Right vmPFC - Control | ↑ Baseline concentration in absence of social cues |
| Right vmPFC - Reputation | ↓ Oxyhaemoglobin concentration compared to baseline, suggesting valuation influenced by reputation |
| Right vmPFC - Face ID and Reputation | ↓ Oxyhaemoglobin concentration compared to baseline, indicating decreased valuation with complex social information |

*Note.* Hemodynamic responses in the ventromedial prefrontal cortex (vmPFC) across different groups. The table summarises the effects of group conditions on oxyhaemoglobin concentration.

## 5.3 Discussion

The goal of this study was to understand the role of dlPFC and vmPFC on the processing of different types of contextual information in cooperative decision-making. Significant effects were observed in both the dorsolateral prefrontal cortex (dlPFC) and the ventromedial prefrontal cortex (vmPFC) across different experimental groups, reflecting changes in oxyhaemoglobin concentration that provide insights into the neural mechanisms underlying social judgement and decision-making. These findings enhance our understanding of how cognitive control, value attribution, and social evaluation interact during cooperative behaviour and add to the extensive literature on value-based behaviour.

In the dlPFC, the Control group demonstrated a decrease in baseline oxyhaemoglobin concentration compared to other groups, suggesting reduced activation. This reduction may indicate a lower demand for cognitive engagement when no specific identity or reputational information is provided and is in concordance with previous literature showing that dlPFC is particularly related to working memory processing and cognitive load (Barbey et al., 2013; Degutis et al., 2024). In contrast, the Reputation group exhibited increased oxyhaemoglobin concentration, indicating heightened cognitive engagement compared to baseline. This increase suggests that participants actively processed reputational information, consistent with the dlPFC’s role in integrating social cues and regulating decision-making based on these inputs (Weissman et al., 2008). The combined Face ID and Reputation group showed even higher oxyhaemoglobin levels, reflecting the joint influence of identity and reputation information on cognitive processes. This joint influence appears to require additional cognitive resources due to the complexity involved in evaluating and integrating multiple social cues simultaneously. This finding suggests that integrating both identity and reputation cues requires additional cognitive resources, likely due to the increased complexity of evaluating multiple types of social information simultaneously.

The Face ID group alone also showed an increase in oxyhaemoglobin concentration, though less pronounced compared to the combined condition (Face ID and Reputation Group), suggesting that identity cues alone necessitate cognitive processing but do not engage the dlPFC as strongly as when combined with reputational information. Interestingly, the Right dlPFC exhibited a decrease in oxyhaemoglobin concentration in the combined Face ID and Reputation group compared to the Face ID group, possibly reflecting an attenuation of cognitive load.

This decrease may imply that when identity information is combined with reputation, the cognitive demands associated with processing these cues are reduced, allowing for more efficient evaluation.

In the vmPFC, distinct patterns of oxyhaemoglobin concentration were also observed across the experimental groups, highlighting its role in value attribution. The Control group exhibited increased oxyhaemoglobin concentration, indicating that the vmPFC was engaged in baseline valuation processes without specific modulatory social information. This suggests a generalised assessment of the decision-making context in the absence of explicit identity or reputational cues. This increased activation may reflect a generalised assessment of the decision-making context in the absence of explicit identity or reputational cues, which is in accordance with evidence of the role of vmPFC to prosocial motivation (Lockwood et al., 2024) and its role on subjective value assessment (Levy & Glimcher, 2012). In contrast, the Reputation group showed a decrease in oxyhaemoglobin concentration, this reduction in activation may indicate that, once the participant has more direct information on what to expect from their co-players, their certainty about the decision can lead to an early confidence representation on the vmPFC (Gherman & Philiastides, 2018). The Face ID and Reputation group demonstrated an increase in concentration, influenced by the presence of multiple sources of social information. This increase suggests that the vmPFC is actively integrating both identity and reputational cues to assign value to the cooperative decision, underscoring its role in synthesising complex social information, which is in conformity with a value-based perspective of vmPFC role on social decision making (Pärnamets et al., 2020).

These findings align with the existing literature on the roles of the dorsolateral and ventromedial prefrontal cortices in cooperation and social decision-making. The increased dlPFC activation observed in the Reputation and Face ID and Reputation groups is consistent with its role in integrating social information, managing cognitive load, and facilitating decision-making during cooperative interactions, as suggested by the value-based framework (Carlson & Crockett, 2018; Pärnamets et al., 2020). The decrease observed in the Right dlPFC for the combined Face ID and Reputation group may indicate that when identity information is integrated with reputation, the cognitive load associated with processing these cues is mitigated, resulting in an attenuation of dlPFC activity. This suggests a potential optimization mechanism, where the brain becomes more efficient at processing complex social information when multiple cues are available.

Similarly, the vmPFC findings are consistent with its role in value attribution, and integration of social information to guide behaviour. The increased activation in the Control group suggests that the vmPFC engages in a baseline valuation process when no specific social information is present, possibly reflecting a generalised appraisal of cooperative decisions. The reduced oxyhaemoglobin concentration observed in the Reputation and Face ID and Reputation groups aligns with the vmPFC’s involvement in assessing social value, particularly when complex reputational information is available. This suggests that the vmPFC plays a critical role in integrating social cues and modulating the perceived value of cooperative actions. When reputational information is provided, the vmPFC may downregulate its activity to reflect decreased motivation to engage in cooperation, especially in the presence of negative reputational cues. Conversely, the increased activation in the Face ID and Reputation group highlights the vmPFC’s role in synthesising diverse social inputs to guide decision-making, suggesting that the presence of identity cues can enhance the perceived value of cooperation, even in the context of reputational challenges.

As limitations, our method to study cooperation was not the most ecological, as Public Goods Games (PGGs) can have several limitations, such as oversimplifying real-world social interactions. Additionally, we did not control for facial characteristics, such as facial expressions or racial traits (e.g., nose and lips shapes), as these aspects were beyond the scope of the current study. This limitation could have influenced how participants perceived their co-players, potentially affecting their cooperative behaviour.

Overall, these results provide further evidence for the differential roles of the dlPFC and vmPFC in processing identity and reputation cues during social decision-making. The dlPFC appears to support cognitive control and the integration of social information, particularly when evaluating the reputational aspects of co-players, while the vmPFC contributes to value-based assessments of social interactions, integrating contextual factors to guide cooperative behaviour. These findings are in line with the value-based framework of cooperation, which posits that cooperative behaviour involves both cognitive and social valuation processes, mediated by distinct but interconnected brain regions. The dlPFC’s role in cognitive control and the vmPFC’s role in value attribution work together to form a cohesive system that enables individuals to effectively navigate complex social environments. This system helps balance social norms, reputational concerns, and personal motivations, ultimately facilitating prosocial actions.

